# Modeling the Biomechanical Features Affecting the Metabolic Rate of Walking with a Powered Ankle-Foot Prosthesis

**DOI:** 10.1101/2025.09.25.678391

**Authors:** Mikayla Schneider, Zane Colvin, Alena M. Grabowski, Cara Gonzalez Welker

## Abstract

For individuals with unilateral transtibial amputation, powered ankle-foot prostheses have the potential to reduce the metabolic rate of walking, which could contribute to improvements in mobility and quality of life; however, physiological improvements have not been consistently demonstrated in experimental studies. To improve our understanding of the biomechanical mechanisms that drive metabolic rate outcomes, we used a machine learning approach to model the relationship between multimodal biomechanical factors and the metabolic rate of walking with a powered ankle-foot prosthesis. Our model included 50 features describing spatiotemporal parameters, step-to-step transition work, joint kinematics, muscle activity, ground reaction forces, prosthesis settings, and subject characteristics, and resulted in a pseudo-R^2^ of 0.986. The features with the largest effect on metabolic rate were peak unaffected side ankle inversion angle, leading affected leg positive work during the step-to-step transition, and peak affected knee extension angle. Accumulated local effects plots were used to visualize the direction and magnitude of the relationship between each feature and the metabolic rate of walking. This work furthers our knowledge about the biomechanical and physiological response to powered ankle-foot prosthesis use and could assist in developing new strategies to drive reductions in metabolic rate.

## 1 INTRODUCTION

Individuals with unilateral transtibial amputation (uTTA) who use a passive-elastic prosthesis incur an elevated metabolic rate of walking compared to people without amputation (Gailey et al., 1994). Reductions in the metabolic rate of walking are associated with improved functional outcomes related to mobility, such as faster preferred walking velocity (Herr and Grabowski, 2012), and greater mobility can indirectly be linked with better quality of life (Mohamadtaghi et al., 2016; Wurdeman et al., 2018). Thus, understanding how to reduce the metabolic rate of walking could contribute to better quality of life in people with uTTA. Ankle-foot prostheses that provide power during the stance phase of walking, such as the BiOM (now Ottobock Empower (Duderstadt, Germany)) [Fig. 1], were developed to improve functional outcomes such as metabolic rate for people with uTTA. Since insufficient push-off power from the prosthetic limb contributes to the elevated metabolic rate for individuals with uTTA (Kuo et al., 2005; Houdijk et al., 2009), supplying additional positive mechanical work at the ankle with a powered prosthesis should normalize metabolic rate levels (Collins and Kuo, 2010; Au et al., 2009; Handford and Srinivasan, 2016). However, the results of experimental studies on individuals with uTTA using such prostheses to walk on level ground have been mixed. Some studies report reductions in the metabolic rate of walking (Herr and Grabowski, 2012; Russell Esposito et al., 2016; Colvin, 2024), while others report no significant change or even an increase in the metabolic rate when individuals with uTTA use a powered prosthesis compared to using a passive-elastic prosthesis (Montgomery and Grabowski, 2018; Kim et al., 2021b; Quesada et al., 2016; Welker et al., 2021; Stafford et al., 2025; Gardinier et al., 2018).The mechanisms behind the metabolic rate changes or lack of changes are unknown. Furthermore, differences between studies in device parameters, such as prosthetic foot stiffness and/or prosthesis power, as well as unique individual adaptations, make it challenging to understand the varied physiological responses to powered prosthesis use.

**Figure 1.**
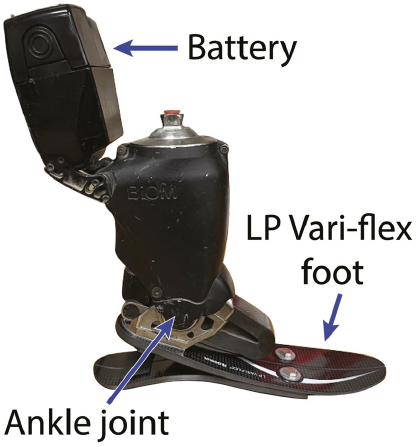
The BiOM T2 battery-powered ankle-foot prosthesis attached in series with the Low Profile (LP) Vari-flex prosthetic foot.

Prior work has revealed some details about the complex interactions between biomechanical factors, device parameters, and the physiological response to walking with a powered prosthesis. For example, Herr and Grabowski (2012) reported that subjects increased trailing affected leg positive work and decreased the magnitude of leading unaffected leg negative work, along with reducing the metabolic rate of walking across a range of speeds using the BiOM compared to a passive-elastic prosthesis. In similar studies of individuals with uTTA using the BiOM by both Mancinelli et al. (2011) and Russell Esposito et al. (2016), peak affected ankle plantarflexion angle and power increased in tandem with reductions in metabolic rate. Regarding muscle activity, Kim et al. (2021a) found no significant correlations between metabolic rate changes and lower-limb muscle activity with the BiOM compared to a passive-elastic prosthesis while walking at a fixed speed normalized to leg length. Differing results have also emerged surrounding prosthesis power magnitude; Ingraham et al. (2018) and Colvin (2024) found that higher settings of BiOM power significantly reduced metabolic rate compared to lower levels and/or passive-elastic prosthesis use, but Quesada et al. (2016) reported no significant differences in metabolic rate for seven conditions with varying peak prosthesis power. While these results have provided some useful insights into the complex response to powered prosthesis use, we still do not understand the full picture of the underlying mechanisms driving changes in metabolic rate.

Of note, much of the previous work relating biomechanical data to metabolic rate using powered prostheses has employed linear models or correlation analyses, which may not capture the full complexity of these relationships. Additionally, given that biomechanical data are highly interrelated, modeling the interactions between multiple data modalities, such as joint angles, muscle activity, and ground reaction forces, with metabolic rate could provide valuable information. However, these data are not always collected simultaneously, and traditional statistical testing cannot handle interaction effects of dozens of biomechanical data features (Quesada et al., 2016). Therefore, an approach that can model numerous nonlinear interactions between multimodal biomechanical data and the metabolic rate of walking could provide valuable insights.

Machine learning models have been used to analyze high-volume, heterogeneous data in the field of biomechanics (Halilaj et al., 2018). Specifically, Bayesian Additive Regression Tree (BART) models have recently been used to analyze the relationships between the metabolic rate of walking and various gait factors for people with cerebral palsy (Gill et al., 2023; Kuska and Steele, 2024; DeVol et al., 2024), including in response to exoskeleton training protocols (Conner et al., 2023). A BART model consists of a sum-of-decision-trees structure that has strong predictive capabilities and incorporates interaction effects of multivariate inputs. Since decision trees are prone to overfitting, the model is regularized by a prior probability distribution, which limits how much each tree can contribute to the overall prediction and results in low levels of bias and variance. Feature selection can also aid in creating statistical models that are stable and not overly sensitive to specific input variables (Luo and Daniels, 2024). Previous work with BART models leveraged accumulated local effects (ALE) plots to compute and visualize the direct effects of each predictor variable on the outcome variable (Apley and Zhu, 2020). By isolating the effect of each biomechanical variable on metabolic rate, ALE plots can help explain which biomechanical variables may be the driving mechanisms behind physiological outcomes.

In order to elucidate the complex interactions while walking with a powered prosthesis, we developed a BART model of the biomechanical features affecting metabolic rate. We used data from 13 individuals with unilateral transtibial amputation walking at 1.25 m/s on level ground using four stiffness categories of passive-elastic prosthetic feet without and with the BiOM stance-phase powered prosthesis at three battery power settings; a total of 16 prosthetic configurations and 196 trials. Multimodal biomechanical data were transformed into predictor features within categories of individual leg step-to-step transition work, lower limb joint angles and moments, leg muscle activity, ground reaction forces (GRFs), and spatiotemporal parameters. Individual features within these categories of data have been previously demonstrated to relate to metabolic rate (Herr and Grabowski, 2012; Russell Esposito et al., 2016; Kim et al., 2021a; Donelan et al., 2001; Grabowski and D’Andrea, 2013), and the use of multiple prosthesis power and stiffness configurations provided a range of biomechanical features and metabolic rates.

We sought to determine and characterize the features with the largest influence on the net metabolic rate of walking in people with uTTA using a passive-elastic and powered prosthesis. We hypothesized that leading unaffected leg negative work, trailing affected leg positive work, and peak affected side plantarflexion angle would be the most important features in terms of magnitude of effect on net metabolic rate, given their previously established correlations with metabolic rate (Herr and Grabowski, 2012; Russell Esposito et al., 2016; Colvin, 2024; Mancinelli et al., 2011).

## 2 MATERIALS AND METHODS

### 2.1 Data Collection

The data for this model were collected from thirteen subjects (3 female/10 male, 41.5 ± 8.5 years, 77.4 ± 15.2 kg) with uTTA. Full details of the data collection are provided by Tacca et al. (2024), and an overview of the protocol is briefly described here [Fig. 2].

**Figure 2.**
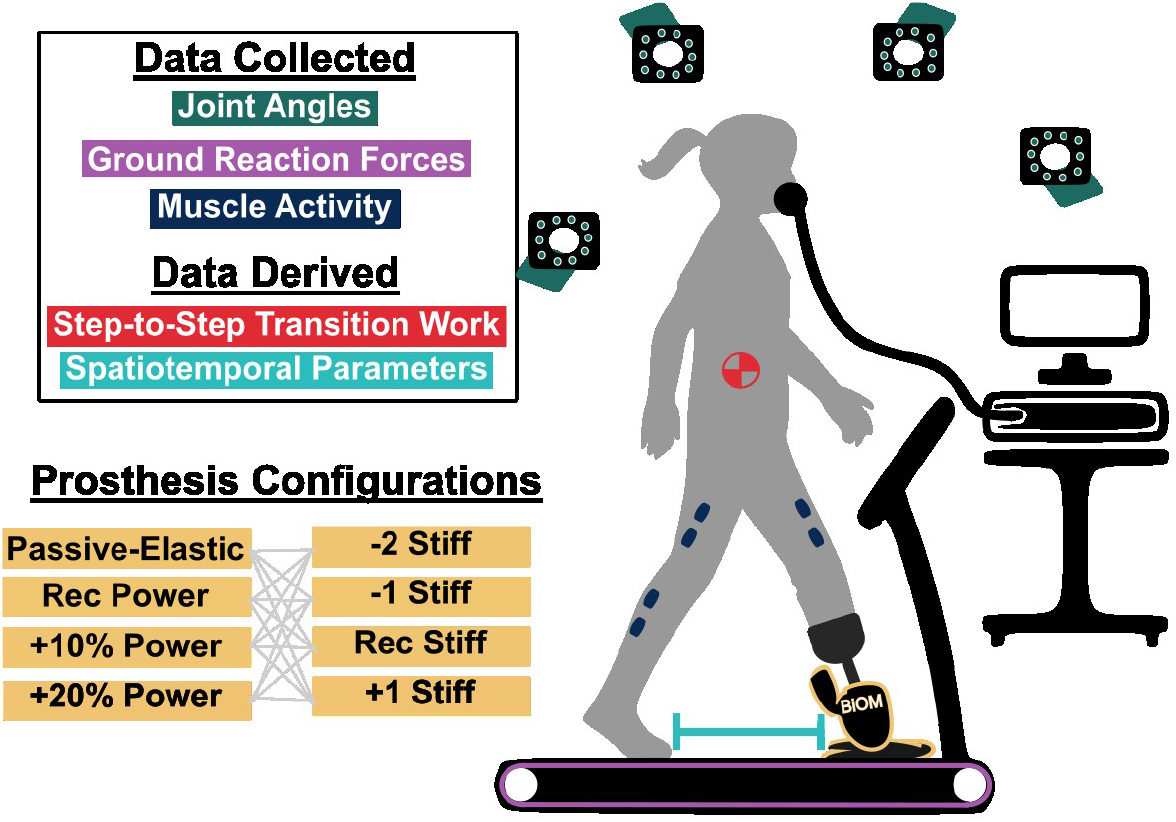
Depiction of the experimental data collection setup. For each trial, motion capture (via cameras and retroreflective markers), ground reaction force (via instrumented treadmill), and muscle activity data (via electrodes) were collected for the lower limbs. From these data, individual leg step-to-step transition work (required to move the center of mass) and spatiotemporal parameters, such as step length, were computed. 16 different prosthesis configurations were tested. The Ossur Low Profile (LP) Vari-flex prosthetic foot at 4 different stiffnesses (Recommended, two levels less stiff, one level less stiff, and one level more stiff) was used along with the BiOM prosthesis at 3 power settings (Recommended, ten percent greater power, and twenty percent greater power). For the “Passive-Elastic” configuration, just the LP Vari-flex foot was used.

In the protocol, each study participant was aligned with a low-profile (LP) Vari-Flex prosthetic foot (Ossur, Reykjavik, Iceland) and the LP Vari-Flex prosthetic foot was then attached to the BiOM by a certified prosthetist. The recommended prosthetic foot stiffness category was defined by the manufacturer based on subject mass and assuming a moderate activity level. Then, participants completed a tuning session in which their recommended power settings with the BiOM were determined. These settings were iteratively set when their prosthetic ankle biomechanics fell within two standard deviations of mean values for able-bodied individuals based on motion capture and GRFs (Jeffers et al., 2015; Jeffers and Grabowski, 2017).

After the tuning session, each subject completed three experimental sessions on different days at the same time of day using 16 total unique prosthetic configurations. The prosthetic foot stiffness categories were: (1) the recommended stiffness category (Rec Stiff), (2) one category stiffer than recommended (+1 Stiff), (3) one category less stiff than recommended (−1 Stiff), and (4) two categories less stiff than recommended (−2 Stiff). The prosthetic battery power settings were: (1) passive-elastic foot only (No Power), (2) recommended power setting (Rec Power), (3) ten percent greater than recommended (+10% Power), and (4) twenty percent greater than recommended (+20% Power). Only power magnitude, not timing, differed between settings.

To collect kinematic data, reflective markers were placed on each subject’s lower limb joint centers (or the equivalent location on the BiOM) and in clusters bilaterally on the foot, shank, thigh, and pelvis segments for optical motion capture. To collect muscle activity, bipolar electrodes (Noraxon, Scottsdale, AZ) were placed on the skin above the biceps femoris long head, gluteus maximus, vastus lateralis, and rectus femoris of both legs, as well as on the lateral gastrocnemius, soleus, and tibialis anterior of the unaffected leg to record surface electromyography (EMG) (Cram et al., 1998). Each subject completed an “EMG baseline trial” at the start of each session, in which they used the recommended prosthetic foot stiffness without the BiOM and walked at 1.25 m/s while EMG data were recorded. This trial was manually checked for electrode failure and reasonable EMG values.

Resting metabolic rate was measured using indirect calorimetry (Parvo Medics TrueOne 2400, Salt Lake City, UT) while subjects stood quietly for five minutes at the start of each session. Then, metabolic rates for each prosthetic configuration were measured while subjects walked on a level split-belt instrumented treadmill (Bertec, Columbus, Ohio) for five minutes at 1.25 m/s. Marker trajectories (100 Hz) (Nexus, Vicon, Centennial, CO), ground reaction forces (1000 Hz), and surface EMG (1000 Hz) were recorded for 30 seconds between minutes 3 and 4 of each walking trial.

### 2.2 Data Processing

To obtain joint angles and moments, the 3D marker positions and ground reaction forces were first low-pass filtered with a fourth-order Butterworth filter with a 7-Hz cut-off frequency (Tacca, 2023). Then, inverse kinematics and dynamics (Visual 3D, C-Motion, Boyds, MD) were applied on a rigid segment model to calculate ankle, knee, and hip joint angles and moments from the marker trajectories.

To obtain muscle activity, the EMG data were demeaned, high-pass filtered with a second-order Butterworth filter at a cut-off frequency of 20 Hz, rectified, and low-pass filtered with a second-order Butterworth filter at a cut-off frequency of 6 Hz. All EMG data were normalized to the average maximum value from the EMG baseline trial for a given muscle, subject, and experimental session day.

All data were segmented into strides from subsequent heel strikes of the same foot, which were identified from vertical GRFs crossing a threshold of 50 N (Rogers-Bradley et al., 2024). Within each subject and variable, mean values from individual strides and time-normalized points within each stride were checked for outliers, defined by 3 standard deviations from the mean. Ankle rotation angles (in the transverse plane) were excluded due to over 30% of the values being removed as outliers. Overall, with the remaining data, less than 5% of strides were removed as outliers.

Spatiotemporal parameters and individual leg step-to-step transition work (Donelan et al., 2001) were calculated from the GRF and 3D marker data. Stride length was calculated as the product of treadmill speed (1.25 m/s) and stride time between consecutive heel strikes on a given side. Stance time was determined as the time between heel strike and toe off on a given side. Step length was calculated as the fore-aft distance between contralateral heel markers. If the affected leg was trailing and the unaffected leg was leading, that step length was labeled as affected-to-unaffected. The same convention applied for the unaffected-to-affected step length. Step width was calculated as the average medio-lateral distance between contralateral heel markers at the instant of leading leg heel strike. Step-to-step transition work was calculated by integrating the external mechanical power, the result of the dot product between the GRF vector and the overall center of mass velocity vector, over periods of double support (Tacca et al., 2024). For each leg, negative and positive individual leg work were found by integrating the negative and positive portions of the external mechanical power, respectively (Donelan et al., 2001).

To account for subject anthropometric differences, stride length, step length, and step width were normalized to subject height. Step-to-step transition work and GRFs were normalized to body mass (including the prosthesis).

Average metabolic rate in W/kg was calculated from the rates of oxygen consumption and carbon dioxide production from the last two minutes of each trial (Brockway, 1987). Then, gross resting metabolic rate was subtracted from each walking trial to obtain net metabolic rate, normalized by body mass (including the prosthesis).

### 2.3 Initial Features

For our initial feature set, we selected values that distinguished individual subjects and trials. We included anthropometric characteristics of age, mass, height, sex, and prosthesis shoe size, which ranged from 22 to 30 cm. We included these participant characteristics in our model as covariates, but they were not the focus of this analysis, which is on the biomechanical and prosthesis features that could be changed, and therefore will not be discussed in the results. From the prosthesis characteristics, we included the researcher-defined power settings and stiffness categories as features (e.g. Rec Power, +1 Stiff, etc.).

Next, we transformed time-series experimental data in order to include them in the BART model, which takes scalar inputs. To characterize muscle activity, we used the integrated value of EMG (iEMG) over a stride, which has been analyzed previously (Kim et al., 2021a; Colvin et al., 2022).

For the biomechanical features, we chose to focus on the stance phase because prior work has shown that the mechanical work required in step-to-step transitions, which occurs during periods of double support, has a significant effect on metabolic rate. Additionally, prosthesis power is only applied during the stance phase. For the joint angles, joint moments, and horizontal and medio-lateral ground reaction forces, the peak values across the stance phase in both directions (e.g. knee flexion and extension) were included as features for each axis and joint, since peak values are the focus of previous literature related to metabolic rates (Mancinelli et al., 2011; Grabowski and D’Andrea, 2013). For the vertical ground reaction forces, we calculated and included values from the first and second peaks. Peak affected side prosthetic ankle inversion and eversion were not included in the model, since this movement is limited with a 1-DoF prosthesis.

If there were any missing features for a given trial, less than 1% on average for all features and trials, we used a random forest algorithm to fill in the missing features based on the rest of the data (Stekhoven and Bühlmann, 2012). This random forest imputation approach preserves as much of the experimental data as possible and has been previously shown to accurately impute missing data values on mixed-type datasets (Stekhoven and Bühlmann, 2012). Only three trials were missing more than 10% of features (21%, 13%, and 11%), and no individual feature was missing more than 5% of values. We also normalized the joint angles and moments, iEMG, GRFs, spatiotemporal parameters, step-to-step transition work, and metabolic rate as z-scores (Conner et al., 2023) relative to the -1 stiffness and no power condition for each subject, which was considered our baseline.

### 2.4 Feature Selection

To ensure robustness in our model, we first narrowed down features by removing those that may be potentially redundant from our initial feature set. From preliminary testing on features to include in the model, we found that peak joint moments in addition to peak joint angles did not improve model performance, and thus we removed peak joint moments from our feature set, leaving us with 68 features remaining. We also evaluated the Pearson correlation coefficient between features to examine redundancy and removed one feature from each pair with a Pearson’s R *>* 0.8: ankle inversion and eversion, hip internal and external rotation, and hip abduction and adduction, on both affected and unaffected sides.

After removing peak joint moments and redundant features, the second stage of feature selection was accomplished using a validated metric that reflects feature importance. For BART models, the importance of each feature in predicting the outcome can be quantified by the Variable Inclusion Proportion (VIP), which is the proportion of times a feature is used as the splitting rule for a tree (Bleich et al., 2014) [Fig. 3A].

**Figure 3.**
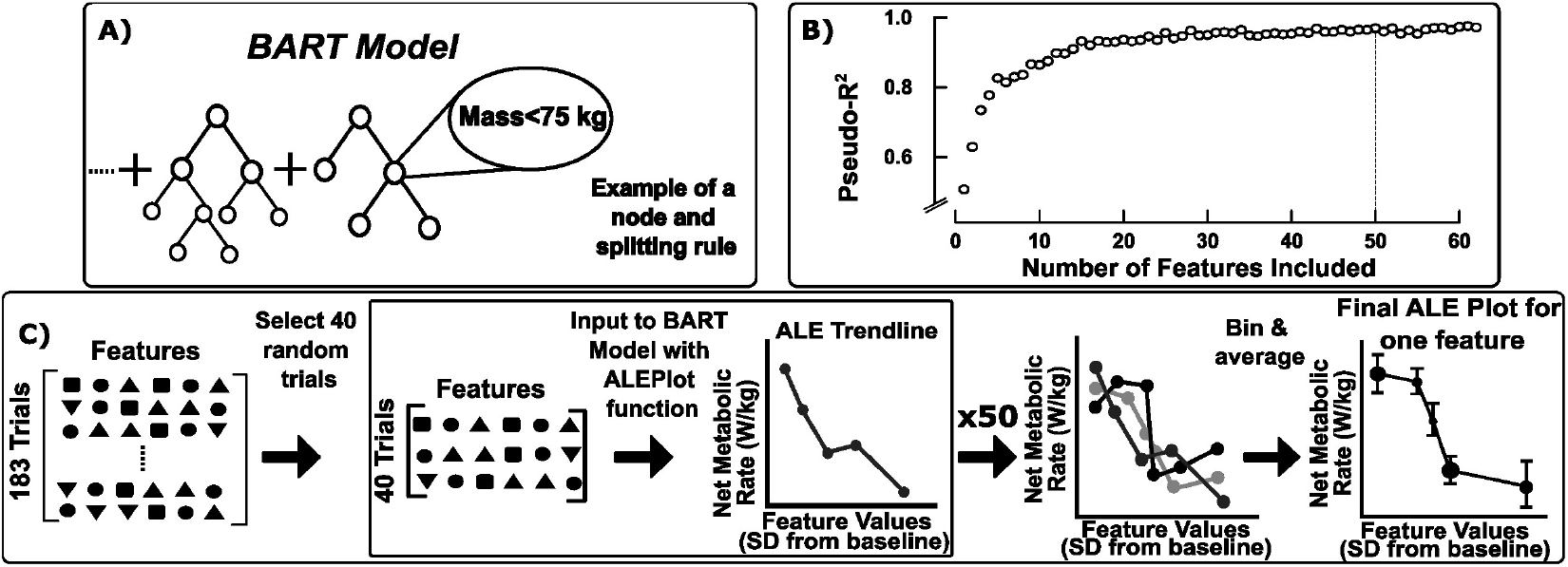
**A)** A high-level visualization of the Bayesian Additive Regression Tree (BART) model structure, with one example of a splitting rule at a node. **B)** Plot of the pseudo-R^2^ values for each of the 62 BART models created to reduce the number of features. The features were ranked according to their Variable Inclusion Proportion (VIP) score and each BART model included increasing numbers of features. The BART model with 50 features yielded a pseudo-R^2^ within 1% of the maximum possible pseudo-R^2^. **C)** The bootstrapping process to create Accumulated Local Effects (ALE) Plots for each feature. 40 trials of all features are randomly selected from the full dataset. This sample is input to the ALEPlot function in Rstudio, which uses the trained BART model to produce an ALE trendline. This process is repeated 50 times to create 50 trendlines from subsets of the full dataset. The trendlines are then binned with even spacing into 10 bins, and the average and standard deviation for each bin is displayed as the final ALE plot.

First, we evaluated the VIP of all remaining 62 features with the *permute*.*vs* (Permutation based Variable Selection) function in R-Studio (Kapelner and Bleich, 2016), which averages the VIP of each feature over 20 distinct BART models (“VIP-test BART models”). Then, we constructed a set of BART models (“feature-test BART models”) with an increasing number of features in order of VIP (i.e. the first model contained one feature, the feature with the highest VIP, and the last model contained all features). We evaluated performance of each feature-test BART model with the pseudo-R^2^ value. Finally, we identified the smallest number of features that resulted in a feature-test BART model with a pseudo-R^2^ within 1% of the maximum pseudo-R^2^ achieved across all feature-test BART models [Fig. 3B]. From this procedure, the first 50 features with the largest VIP from the VIP-test models (Table I) were selected for use in the final model.

**Table 1.** The 50 final model features.

| Category | Individual Features |
| --- | --- |
| Subject | Mass, Height, Sex, & Prosthesis Shoe Size |
| Prosthesis | Power Setting & Stiffness Category |
| Spatiotemporal Parameters | Unaffected & Affected Stance Time, Average Step Width, Unaffected-to-Affected Step Length |
| Joint Kinematics | Peak Unaffected Ankle Inversion, Affected & Unaffected Plantarflexion & Dorsiflexion, Affected & Unaffected Knee Extension & Flexion, Affected & Unaffected Hip Extension, Flexion & Internal Rotation, Affected Hip Abduction |
| Muscle Activity | Integrated EMG of Unaffected Biceps Femoris, Rectus Femoris, Gluteus Maximus, Tibialis Anterior, Soleus, & Lateral Gastrocnemius, Affected Biceps Femoris & Rectus Femoris |
| Ground Reaction Forces | Peak Affected Lateral & 2nd Peak Affected Vertical, Affected & Unaffected Medial & Horizontal Propulsive, 1st and 2nd Peak Unaffected Vertical, Unaffected Braking GRF |
| Step-to-Step Transition Work | Leading Affected & Unaffected Leg Positive & Negative, Trailing Affected & Unaffected Leg Positive, Trailing Affected Leg Negative Work |

### 2.5 Final BART Model Development

We developed the final BART model with the *bartMachineCV* package in RStudio [Fig. 3] (Kapelner and Bleich, 2016). With this package, the model’s hyperparameters, variables that control how the model is trained, were automatically tuned through k-fold cross-validation over a grid of options. This function created multiple BART models, each with a unique combination of four hyperparameters, such as the number of trees. The *bartMachineCV* function then selected the option with the lowest average out-of-sample root mean square error (RMSE), which is the RMSE evaluated on each of the randomly selected k groups of held-out test data.

After development of the final BART model, we leveraged accumulated local effects (ALE) plots to interpret the effects of each predictor variable. Accumulated local effects are the integral of the average change in the outcome variable because of a small change in a given window of the predictor variable value. ALE plots are unbiased for highly correlated features, unlike similar techniques such as partial dependence plots (Apley and Zhu, 2020). For our model, the ALE plots display the average change in net metabolic rate that results from a change in the predictor variable.

ALE plots were generated for each feature through a bootstrapping process [Fig. 3C] in order to obtain a distribution of the sample (Henderson, 2005). In this process, 40 walking trials were first randomly sampled from the 182 trials remaining after the z-score normalization. These 40 trials were input to the *ALEPlot* function (Apley, 2018) with the trained BART model to generate the partial effects on net metabolic rate for each feature, which produces an ALE plot with heterogeneous X and Y spacing. This process is then repeated 50 times. The resulting ALE plots are binned into 10 evenly spaced increments as a method of averaging and smoothing. The final values are plotted as an average and standard deviation value, where the size of the data point denotes the number of values in that bin. The total effect size of each feature was then calculated as the difference between the 90th and 10th percentile of the ALE plot values.

To analyze the relationship between features, we calculated Pearson correlation coefficients using the normalized features as they were input to the BART model. Pearson correlation coefficients range from -1 (perfect negative linear relationship) to 1 (perfect positive linear relationship). We interpreted coefficient magnitudes of 0.4-0.69 as moderate, 0.7-0.89 strong, and 0.9-0.99 very strong (Schober et al., 2018).

## 3 RESULTS

5-fold cross validation was used to train the final model, which had a pseudo-R^2^ value of 0.986 and an in-sample normalized RMSE of 5.88%. Fig. 4 displays the features (excluding subject characteristics) with an effect size greater than the model’s RMSE. Individual model features had effect sizes that ranged from 0.18 to 0.65 W/kg (5.9 to 21% of the average net metabolic rate). The top features were distributed across categories of step-to-step transition work, muscle activity, joint angles, spatiotemporal parameters, ground reaction forces, and prosthesis configuration features for both legs.

**Figure 4.**
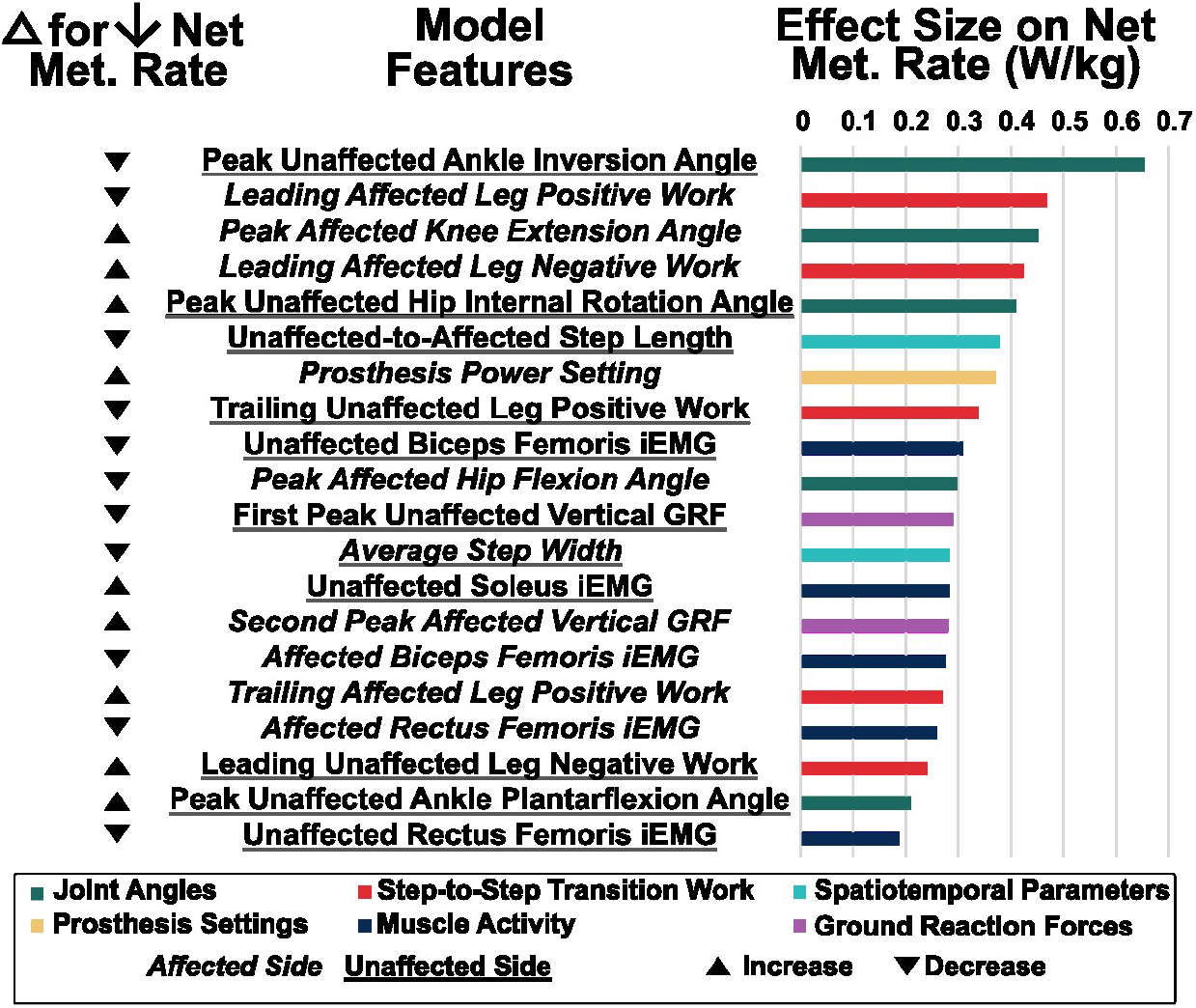
Effect sizes on net metabolic rate of the features with an effect size greater than the model’s RMSE. The leftmost column displays the general direction of change (increase or decrease) to reduce the net metabolic rate of each feature according to its ALE plot.

First, we generated the ALE plots and calculated the effect size for the features considered in our hypothesis, as well as prosthesis power setting, since these features have been extensively examined in prior work [Fig. 5]. Trailing affected leg positive work (effect size = 0.27 W/kg), leading unaffected leg negative work (effect size = 0.24 W/kg), and peak affected ankle plantarflexion angle (effect size = 0.09 W/kg) each had roughly nonlinear negative correlations with net metabolic rate. Prosthesis power settings ten and twenty percent higher than the recommended prosthesis power setting (effect size = 0.37 W/kg) correlated with lower net metabolic rate.

**Figure 5.**
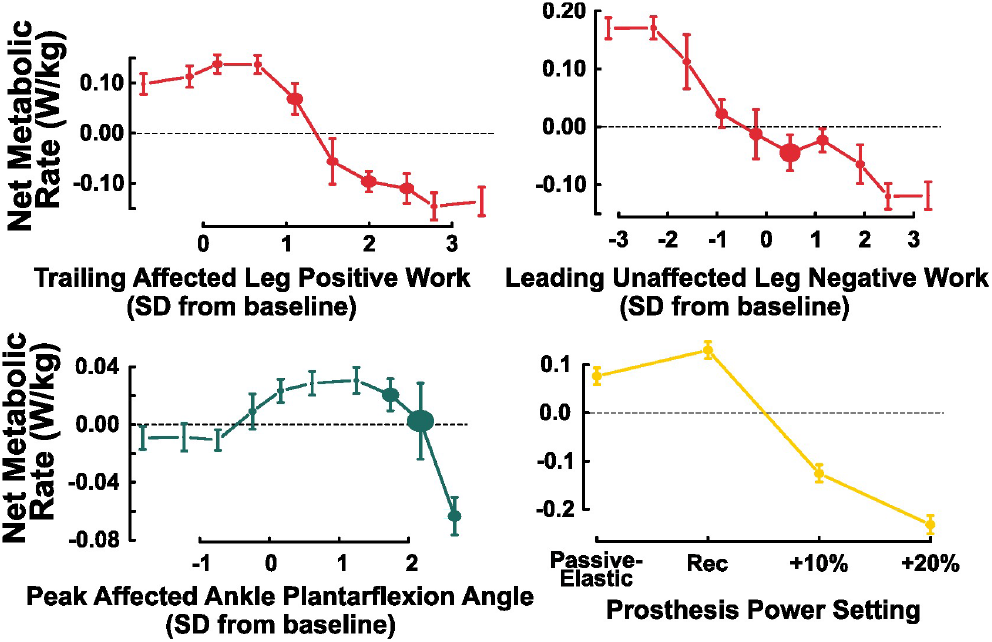
ALE plots of biomechanical features that have been studied in previous work (from left to right, top to bottom): trailing affected leg positive work, magnitude of leading unaffected leg negative work, peak affected ankle plantarflexion angle, and prosthesis power setting. Our model results demonstrate similar trends as prior results. The baseline condition is that of the passive-elastic prosthesis with the -1 stiffness category.

We also generated the ALE plots for the next eight features with the largest effect sizes that have not been assessed experimentally in prior work [Fig. 6]. Peak unaffected ankle inversion angles during the stance phase (effect size = 0.65 W/kg), leading affected leg positive work (effect size = 0.47 W/kg), unaffected-to-affected step length (effect size = 0.38 W/kg), trailing unaffected leg positive work during the step-to-step transition (effect size = 0.34 W/kg), and peak affected hip flexion angles during the stance phase (effect size = 0.30 W/kg) each had nonlinear negative correlations with net metabolic rate. The magnitude of peak affected knee extension angle (effect size = 0.45 W/kg), leading affected leg negative work during the step-to-step transition (effect size = 0.42 W/kg), and peak unaffected hip internal rotation angle (effect size = 0.41 W/kg) each had a nonlinear positive correlation with net metabolic rate. Values of unaffected biceps femoris iEMG (effect size = 0.31 W/kg) that were either greater than or one standard deviation less than the baseline were correlated with increased net metabolic rate.

**Figure 6.**
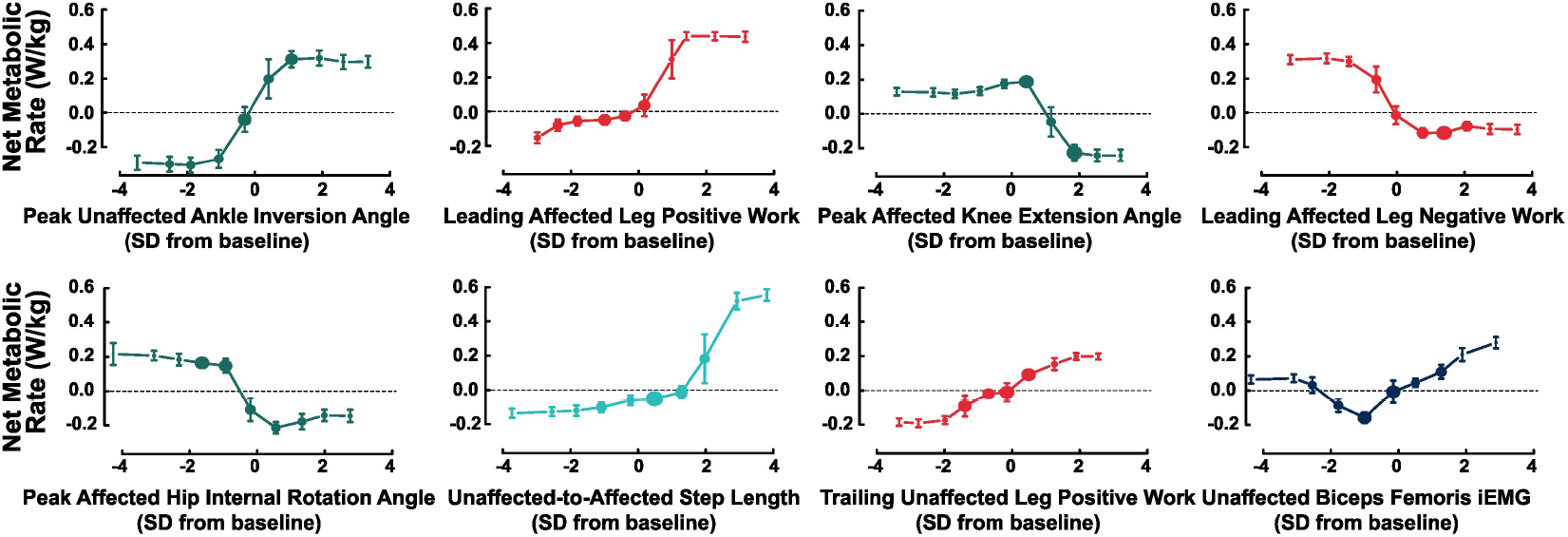
ALE plots of the remaining 8 features that had the largest effect. The plots are displayed from left to right, then top to bottom in order of total effect magnitude. The plot colors correspond to the feature type (e.g. dark blue for muscle activity). All features represent the magnitude of the feature value (e.g. the magnitude of leading affected leg negative work). The baseline condition is that of the passive-elastic prosthesis with the -1 stiffness category.

## 4 DISCUSSION

We used a machine learning model to identify the most important features relating to net metabolic rate for individuals with uTTA walking with a powered ankle-foot prosthesis on level ground. From our results, we reject our hypothesis that the previously analyzed features of trailing affected leg positive work, leading unaffected leg negative work, and peak affected ankle plantarflexion angle would have the largest effect on net metabolic rate, as these features were ranked 16, 18, and 39 out of the 46 non-subject-characteristic features, respectively, in terms of effect size.

Although the trailing affected leg positive work and leading unaffected leg negative work features have relatively small effect sizes and low rank, other step-to-step transition work features, including leading affected leg positive work (rank 2/46), leading affected leg negative work (rank 4/46), and trailing unaffected leg positive work (rank 8/46) have a large impact on metabolic rate in our model. This aligns with prior work reporting that the elevated metabolic rate during walking for individuals with amputation is partly attributable to the elevated costs of transitioning the center of mass (Houdijk et al., 2009). The importance of leading unaffected leg negative work may be reflected by the first peak of unaffected leg vertical GRF (rank 11/46), since a moderate linear correlation (Pearson’s R = 0.65) exists between the two features. Similarly, the importance of prosthesis power setting (rank 7/46) may reflect the importance of trailing affected leg positive work (Pearson’s R = 0.77) and peak affected ankle plantarflexion angle (Pearson’s R = 0.57). Our model demonstrates that these previously determined features are still indirectly important to metabolic rate, but are not the most influential.

The finding that prosthesis power setting does not have the largest effect on metabolic rate (rank 7/46) is particularly surprising given the amount of work focused on applying biologically-inspired power to prosthetics and exoskeletons (Quesada et al., 2016; Colvin et al., 2022; Ingraham et al., 2018; Mooney et al., 2014). Our model trend for prosthesis power setting matches previous work demonstrating that a BiOM prosthesis power setting greater than the recommended correlates with reduced metabolic rate (Ingraham et al., 2018; Colvin, 2024); however, its effect size is smaller than that of six biomechanical features, indicating the importance of user adaptation to this power. It is important to note that one prosthesis power condition included in the model was no power (with only a passive-elastic foot), so this result reflects the magnitude difference not just between power settings, but between powered and passive-elastic prostheses. In future work, it may be advantageous to place a greater focus on how the prosthesis users are adapting to the applied power and develop strategies that incorporate information about additional biomechanical feature changes to take full advantage of the prosthesis power.

While the magnitude of their impact may have been smaller than expected, the model-predicted trends for trailing affected leg positive work, leading unaffected leg negative work, and peak affected ankle plantarflexion angle agree with prior experimental results. First, our model results match the previously reported correlations between lower metabolic rate and greater trailing affected leg positive work (Herr and Grabowski, 2012; Russell Esposito et al., 2016). Second, in alignment with Russell Esposito et al. (2016), our model demonstrated correlations between lower metabolic rate and increased magnitudes of leading unaffected leg negative work. Third, the correlation between peak affected plantarflexion angle and reduced net metabolic rate, as found by Mancinelli et al. (2011) and Russell Esposito et al. (2016), is broadly seen in our model, with slight discrepancies. Specifically, the model predicts that the lowest net metabolic rate occurs at the highest peak affected ankle plantarflexion angles, with a nonlinear trend at lower ankle plantarflexion angles (Fig. 5). These slight discrepancies from prior work could be attributed to the fact that we calculated peak plantarflexion angle during stance phase prior to toe-off (instead of over the entire gait cycle as in prior work), so our model lacked information about the swing phase. Overall, the alignment between previous identified trends and our model trends for the three features discussed here, as well as prosthesis power setting, give us confidence in our model.

Unsurprisingly, the top features in our model were a mix of joint angles, step-to-step transition work, spatiotemporal parameters, prosthesis settings, muscle activity, and ground reaction forces for the affected and unaffected legs. This suggests that the physiological response to using a powered prosthesis is a complex process based on multiple factors, including an individual’s biomechanical adaptation to the prosthesis. Walking with an assistive device requires complex coordination, as multiple biomechanical features can change in tandem. The mechanisms behind metabolic rate changes with exoskeleton use are similarly complex; one study by Poggensee and Collins (2024) reports that exoskeleton users who reduce metabolic rate demonstrate opposing trends in changes in joint work rate, muscle activity and kinematics at the assisted vs. the unassisted joints. Our results showing that a diversity of feature types strongly affect metabolic rate support the use of multimodal data; restricting our model to just one type of data likely would have limited its performance. Given the many different important features, we identified overarching themes based on prior work on prostheses and metabolic rate that may explain how these features could work together to reduce metabolic rate: equal magnitudes of trailing leg positive work and leading leg negative work, similar magnitudes of both positive and negative step-to-step transition work for the affected and unaffected legs, and frontal plane stability.

Equal magnitudes of trailing leg positive work and leading leg negative work have been theorized to minimize net metabolic rate in able-bodied walking (Kuo et al., 2005). This theory may explain our model’s finding that greater magnitudes of leading unaffected leg negative work correlate with reduced metabolic rate. While our results align with those from Russell Esposito et al. (2016), Herr and Grabowski (2012) reported the opposite trend, leaving the relationship between leading leg negative work and metabolic rate unclear. When using an inverted pendulum model to represent each leg in theoretical simulations of steady-state walking, the transition from one leg to the next is most economical with equal magnitudes of trailing leg positive work and leading leg negative work during double support; any imbalance requires additional work in single support, which would increase the net metabolic rate Kuo et al. (2005). Therefore, our model trends correlating reduced net metabolic rate with increased magnitudes of leading unaffected leg negative work and increased magnitudes of trailing affected leg positive work could make sense in the context of this theoretical model.

Similar magnitudes of positive step-to-step transition work for the affected and unaffected legs has been used as a metric of evaluating powered prosthesis performance (Montgomery and Grabowski, 2018) and may be related to reduced metabolic rate (Adamczyk and Kuo, 2015). Individuals with uTTA who use a passive-elastic prosthesis generate more positive work during push-off and mid-stance with their unaffected side compared to their affected side; this contrasts with those without amputation, who generate equal amounts of positive work with both legs (Adamczyk and Kuo, 2015). Our model trends showing correlations between reduced metabolic rate and less trailing unaffected leg positive work, less leading affected leg positive work, and greater trailing affected leg positive work reflect an offloading of positive work to the battery-powered prosthesis. Additionally, our model results suggesting that shorter unaffected-to-affected step lengths correlate with reduced net metabolic rate may align with the reduced push-off work from the trailing unaffected leg, along with the reduced biceps femoris muscle activity through its attachment at the hip. Future work would be needed to directly test the impact of similarities in magnitude in positive work between the affected and unaffected legs, but these combined features suggest that equal amounts of individual leg positive work may be associated with reduced net metabolic rate.

Similar magnitudes of negative step-to-step transition work for the affected and unaffected legs may also be a goal for powered prosthesis users to reduce metabolic rate. With a passive-elastic prosthesis, the magnitude of leading affected leg negative work is reduced compared to that of the unaffected leg, which may be due to a reliance on trailing unaffected leg positive work to propel the center of mass forward and maintain walking speed (Adamczyk and Kuo, 2015). Our model indicates that greater magnitudes of leading affected leg negative work are associated with reduced metabolic rate and points to a number of kinematic changes that could synergistically result in increased leading affected leg negative work. For example, leading affected leg negative work could be aided by an extended leg position during early stance, characterized by reduced peak hip flexion angle and increased peak knee extension angles, which our model associates with reduced metabolic rate.

The importance of reduced peak unaffected ankle inversion angle during stance (rank 1/46), increased peak unaffected hip internal rotation angle during stance (rank 5/46), and reduced step width (rank 12/46) in reducing metabolic rate likely indicate that maintaining frontal plane stability is a key component influencing net metabolic rate. Prior work by Kim and Collins has demonstrated the importance of frontal plane stability for individuals with uTTA using a powered prosthesis, who experienced a lower metabolic rate of walking with a controller that assisted with stability by providing ankle inversion/eversion torque compared to a zero-gain controller (Kim and Collins, 2017). More generally, the relationship between step width and metabolic rate has been established by Donelan et al. (2001), who reported that metabolic rate increases at step widths greater than and less than the preferred width. This aligns with our model trend relating reductions in step width from the baseline condition with reductions in net metabolic rate. In addition, greater ankle inversion angles shift the ankle joint center laterally in the frontal plane, leading to greater step widths, which are associated with fall risk (Maki, 1997) and decreased stability control (Molina et al., 2023). Finally, increased hip internal rotation helps to direct the lower limb more medially in the frontal plane. Therefore, maintaining stability in the frontal plane by decreasing peak unaffected ankle inversion angle, increasing peak unaffected hip internal rotation angle, and reducing step width may be a strategy used to reduce metabolic rate according to our model results.

Our machine learning model provides important insights into the biomechanical features that have the largest effect on the net metabolic rate of walking using a stance-phase powered ankle-foot prosthesis compared to a passive-elastic prosthesis; however, there are limitations to the model. First, the BART model structure requires a scalar value to represent each feature and cannot utilize time-series data. We chose to use peak values for the features of most data (with the exception of integrating muscle activity), but features could also be created with timing onset of peak values, values at specific time points such as heel strike or toe-off, principal component analysis, and other approaches. Additionally, the model is not constrained to obey the musculoskeletal dynamics and physiology of human motion, nor does it assume any causal relationships. Incorporating additional features and/or analyzing these model results within physically-realistic constraints could yield useful insights.

Although these results are promising, further work is required to confirm the validity of the associations between biomechanical features and net metabolic rates for walking. The overarching themes presented– equal magnitudes of trailing leg positive work and leading leg negative work, similar magnitudes of both positive and negative step-to-step transition work for the affected and unaffected legs, and frontal plane stability– were discussed to suggest how certain biomechanical features could be aligned given the inherent interactions between all features. Future studies would be required to verify these themes as approaches to reduce net metabolic rate, since the themes were not explicitly modeled in our work.

In future work, musculoskeletal simulation software such as OpenSim (Seth et al., 2018) could be used to evaluate the identified biomechanical feature changes with dynamically and physiologically-realistic models. Additionally, important features could be experimentally tested with gait retraining protocols; for example, individuals with uTTA who use a BiOM could be given instruction and feedback to decrease their peak unaffected ankle inversion angle, theoretically leading to reductions in metabolic rate. This future work would validate the results of this model and establish the protocol for data-informed insights to drive reductions in net metabolic rate.

## 5 CONCLUSION

The objective of this study was to determine the influence between biomechanical factors on the net metabolic rate of walking with a powered prosthesis compared to a passive-elastic prosthesis. Using an experimental dataset of individuals with uTTA walking at 1.25 m/s on level-ground using a passive-elastic prosthetic foot and a BiOM prosthesis at different power settings and stiffness categories, we developed a machine learning model that associated specific biomechanical features with changes in net metabolic rate. We incorporated joint kinematics, peak ground reaction forces, integrated muscle activity, spatiotemporal parameters, subject characteristics, and prosthesis configurations together in a single model to explain changes in net metabolic rate. This model and approach allows us to visualize the individual relationships between each of the biomechanical features with net metabolic rate and to compare their relative effect sizes. The results of the model agree with previously established relationships between net metabolic rate and step-to-step transition work, prosthesis power setting, and affected side peak plantarflexion angle. In addition, the model elucidates further associations with other biomechanical features, including peak unaffected side ankle inversion angle, leading affected leg positive work during the step-to-step transition, and peak affected knee extension angle.

These results provide a framework for understanding the biomechanical underpinnings of users’ metabolic responses to walking using a powered ankle-foot prosthesis. Additionally, the features most strongly associated with net metabolic rates could inform new walking strategies for powered prosthesis users and/or novel prosthesis designs or control schemes. Overall, the results of this research can inform progress in enhancing performance and adoption of powered prostheses that improve the quality of life for individuals with amputation.

## AUTHOR CONTRIBUTIONS

MS: Conceptualization, Formal Analysis, Methodology, Visualization, Writing - original draft, Writing - review & editing; ZC: Investigation, Data Curation, Writing - review & editing; AMG: Funding Acquisition, Resources, Writing - review & editing; CGW: Conceptualization, Supervision, Writing - review & editing

## ACKNOWLEDGMENTS

The authors thank Joshua Tacca for his assistance in data collection.

